# Frequency band coupling with high-frequency activities during seizures shifts from θ band in tonic phase to δ band in clonic phase

**DOI:** 10.1101/2021.10.06.463288

**Authors:** Hiroaki Hashimoto, Hui Ming Khoo, Takufumi Yanagisawa, Naoki Tani, Satoru Oshino, Masayuki Hirata, Haruhiko Kishima

## Abstract

**Objective:** To clarify variations in the relationship between high-frequency activities (HFAs) and low-frequency bands from the tonic to the clonic phase in focal to bilateral tonic-clonic seizures (FBTCS), using phase-amplitude coupling.

**Methods:** This retrospective study enrolled six patients with drug-resistant focal epilepsy who underwent intracranial electrode placement for presurgical invasive electroencephalography at Osaka University Hospital (July 2018–July 2019). We used intracranial electrodes to record seizures in focal epilepsy (11 FBTCS). The magnitude of synchronization index (SIm) and receiver-operating characteristic (ROC) analysis were used to analyze the coupling between HFA amplitude (80–250 Hz) and lower frequencies phase.

**Results:** The θ (4–8 Hz)-HFA SIm peaked in the tonic phase, whereas the δ (2–4 Hz)-HFA SIm peaked in the clonic phase. ROC analysis indicated that the δ-HFA SIm discriminated well the clonic from the tonic phase.

**Conclusions:** The main low-frequency band modulating the HFA shifted from the θ band in the tonic phase to the δ band in the clonic phase.

**Significance:** In FBTCS, low-frequency band coupling with HFA amplitude varies temporally. Especially, the δ band is specific to the clonic phase. These results suggest dynamically neurophysiological changes in the thalamus or basal ganglia throughout FBTCS.

**Highlights:**

- The θ band (4–8 Hz) was mainly coupled with high-frequency activity (HFA) in the tonic phase of focal to bilateral tonic-clonic seizures (FBTCS).
- The δ band (2–4 Hz) was mainly coupled with HFA in the clonic phase of FBTCS.
- The magnitude of the synchronization index related to δ-HFA phase-amplitude coupling discriminated well the clonic from the tonic phase.

## 1. Introduction

Focal to bilateral tonic-clonic seizure (FBTCS) is a new term, revised in 2017 by the International League Against Epilepsy, which was referred previously as secondarily generalized seizure. The FBTCS reflects the propagation pattern of a seizure (Fisher et al., 2017). At initial diagnosis, it is observed in approximately 80% of the patients with focal epilepsies (Forsgren et al., 1996). FBTCS is the most important risk factor for sudden unexpected death in epilepsy (Shorvon and Tomson, 2011) and a poor prognosis factor for epilepsy surgery (Bone et al., 2012). However, antiepileptic drugs to prevent FBTCS remain unknown (Hemery et al., 2014), and it remains unclear why some patients suffer from uncontrolled FBTCS and others do not (He et al., 2020, Sinha et al., 2021). Uncovering the neurophysiological mechanism underlying FBTCS may provide new insights into the development of new treatment strategies.

Diffusion-weighted magnetic resonance imaging (MRI) studies have revealed that patients with drug-resistant temporal lobe epilepsy and FBTCS have widespread abnormalities in whole-brain structural networks (Sinha et al., 2021). In the subcortical region, the thalamus is hyperactivated during FBTCS (Hamandi et al., 2006) and may play an essential role in the propagation of ictal activity to widespread cortical networks (Castro-Alamancos, 1999). A thalamic marker of a functional MRI study has been shown to discriminate between individuals with and without FBTCS (Caciagli et al., 2020). The basal ganglia network is known as a braking system between the cortex and thalamus (Vuong and Devergnas, 2018) and is hyperactive during FBTCS (Blumenfeld et al., 2009). It has been proposed that FBTCS may reflect a broken balance between basal ganglia inhibition and thalamic synchronization (He et al., 2020).

FBTCS has also been investigated using neurophysiological oscillatory activities, such as high-frequency activities (HFAs). Prior to the onset of FBTCS, HFAs inside the seizure onset zone (SOZ) are higher in FBTCS than in focal seizures (Schönberger et al., 2019) and HFAs are observed during intracranial postictal attenuation related to the FBTCS (Bateman et al., 2019). Tonic-clonic phases in FBTCS show significantly higher values of HFAs than the non-motor symptom phase in FBTCS (Hashimoto et al., 2021b). HFA is a key oscillation in epilepsy research (Jirsch et al., 2006, Modur et al., 2012, Zijlmans et al., 2012) and is reported to be related to physiological neural-oscillatory changes (Hashimoto et al., 2017, Hashimoto et al., 2020a, Hashimoto et al., 2021d). As such, it is clinically important to distinguish between physiological and pathological HFAs in epileptic patients (Cimbalnik et al., 2018, Frauscher et al., 2018, Matsumoto et al., 2013).

Some studies have investigated HFAs using the phase-amplitude coupling (PAC) method in combination with lower-frequency activities (Canolty et al., 2006). PAC analysis measures the degree of synchronization between phases with low-frequency oscillations and high-frequency amplitudes (Cohen, 2008). PAC can be physiological (Hashimoto et al., 2021c, Yanagisawa et al., 2012) or pathological. Studies related to pathological PAC have demonstrated that ictal HFA amplitude is coupled with δ (Nariai et al., 2011) or θ (Ibrahim et al., 2014) bands. Our previous studies investigating FBTCS have shown that PAC between infra-slow activities and HFAs precede ictal HFAs (Hashimoto et al., 2020b, 2021a). Moreover, previous studies have mainly focused on ictal PAC or PAC of the seizure onset zone (SOZ), and it remains unclear how PAC dynamically changes throughout FBTCS.

Our previous study showed that, at the beginning of seizures, θ-HFA coupling and ictal HFA power increase, followed by a decrease in θ-HFA coupling; however, δ-HFA coupling increases after θ-HFA coupling increases (Hashimoto et al., 2021b). Another study also reported that the main low-frequency band coupled with HFA shifts from 4–5 Hz (θ band) to 1–2 Hz (δ band) during the seizure (Grigorovsky et al., 2020).

In this study, we hypothesized that, in FBTCS, θ-HFA coupling may occur in the tonic phase and δ-HFA coupling may occur in the clonic phase. Besides the SOZ, we also investigated all contacts that showed significant coupling, including those observed in the propagation zone. Moreover, we used the synchronization index (SI) (Cohen, 2008) to measure the strength of PAC between the HFA amplitude and the θ and δ phases.

## 2. Materials and Methods

### 2.1. Subjects and study setting

In this retrospective study, we enrolled six patients with drug-resistant focal epilepsy who underwent intracranial electrode placement for presurgical invasive electroencephalography (EEG) and who were admitted to Osaka University Hospital between July 2018 and July 2019 (Table 1). Some patients (P1 to P5) in this study were the same patients as those included in our previous study (Hashimoto et al., 2021a, 2021b).

**Table 1.** Clinical profile of the enrolled patients

| Patient number | Sex | Age at surgery (years) | Pathology | Seizure number | Tonic phase (Duration) | Clonic phase (Duration) | Number of total implanted contacts |
| --- | --- | --- | --- | --- | --- | --- | --- |
| P1 | Male | 28 | R MTLE<br>DNT | S1 | Yes (28 s) | Yes (33 s) | 96 |
|  |  |  |  | S2 | Yes (24 s) | Yes (29 s) |  |
|  |  |  |  | S3 | Yes (24 s) | Yes (30 s) |  |
| P2 | Male | 47 | R PLE<br>Glial-cortical tissue<br>with psammoma body-<br>like lesion | S1 | Yes (13 s) | Yes (11 s) | 54 |
| P3 | Male | 20 | L OLE* | S1 | Yes (34 s) | Yes (27 s) | 92 |
| P4 | Female | 15 | L MTLE<br>HS | S1 | Yes (19 s) | Yes (19 s) | 72 |
|  |  |  |  | S2 | Yes (27 s) | Yes (29 s) |  |
| P5 | Male | 15 | R OLE<br>Diffuse glioma | S1 | Yes (16 s) | Yes (17 s) | 62 |
|  |  |  |  | S2 | Yes (22 s) | Yes (17 s) |  |
|  |  |  |  | S3 | Yes (19 s) | Yes (17 s) |  |
| P6 | Male | 24 | L MTLE<br>Cortical dysplasia | S1 | No | Yes (44 s) | 74 |

Notes: The seizure number is the serial number of seizures in each participant. All seizures are both FIAS and FBTCS.
Abbreviations: DNT, dysembryoplastic neuroepithelial tumor; FBTCS, focal to bilateral tonic-clonic seizure; FIAS, focal impaired-awareness seizure; HS, hippocampal sclerosis; L, left; MTLE, mesial temporal lobe epilepsy; OLE, occipital lobe epilepsy; PLE, parietal lobe epilepsy; R, right
\* The electrodes were placed primarily in the temporal lobe because we suspected MTLE based on the pre-surgical examination, and a few electrodes were placed in the occipital lobe. The intracranial EEG study revealed OLE, but the range of locations where the intracranial electrodes were placed was not sufficiently wide to detect the exact seizure onset zone. Thus, we decided not to perform a focal resection surgery.

This study was approved by the Ethics Committee of Osaka University Hospital (Suita, Japan) (approval no. 19193) and was conducted in accordance with the Declaration of Helsinki for experiments involving humans. Informed consent was obtained using the opt-out method from our center’s website. We confirmed that all methods were performed in accordance with the relevant guidelines and regulations.

### 2.2. Intracranial electrodes

Intracranial EEG (iEEG) data were acquired using a combination of subdural grids (10, 20, or 30 contacts), strips (four or six contacts), and depth electrodes (six contacts) (Unique Medical Co. Ltd., Tokyo, Japan), placed using conventional craniotomy. The diameter of each contact was 3 or 5 mm, and the inter-contact distances were 5, 7, or 10 mm for the grid and strip electrodes. The diameter and inter-contact distance were 1.5 mm and 5 mm, respectively. Three-dimensional (3D) brain renderings were created using FreeSurfer (https://surfer.nmr.mgh.harvard.edu) on the preoperative MRI images. Using Brainstorm (http://neuroimage.usc.edu/brainstorm/), post-implantation computerized tomography images were overlaid onto the 3D brain renderings. The location and laterality of the intracranial electrodes were determined by presurgical examinations, such as scalp EEG, MRI, and fluorodeoxyglucose-positron emission tomography. The total number of implanted contacts is listed in Table 1.

### 2.3. Data acquisition and preprocessing

Signals from iEEG were acquired at a sampling rate of 1 kHz and a time constant of 10 s by using a 128-channel digital EEG system (EEG 2000; Nihon Kohden Corporation, Tokyo, Japan). The EEG system was also equipped with video (Video-EEG monitoring). BESA Research 6.0 software (BESA GmbH, Grafelfing, Germany) was used to preprocess the raw signals by using a 60-Hz notch filter with a 2-Hz width to eliminate the alternating current line artifact and a zero-phase low-pass filter at 333 Hz with a 24-dB/oct slope to prevent aliasing. Next, the BESA software exported the data to a text file containing iEEG signals. This was then imported to MATLAB R2020b (MathWorks, Natick, MA, USA), and iEEG signals were digitally re-referenced to the common average of all implanted contacts in each patient. The common average was calculated from the mean of all contacts and subtracted from each value acquired from each contact.

We saved iEEG data every 60 min; therefore, one text file contained one 60-minute signal. We applied a bandpass filter to the entire 60-minute data to prevent edge-effect artifacts.

### 2.4. Tonic and clonic phase

Seizure onset was determined by visual inspection of iEEG signals using low-voltage fast activity (Perucca et al., 2014), disappearance of the background activity (Ikeda et al., 1999), and direct current shifts (Ikeda et al., 1996). All investigated seizures in this study were FBTCS, which contained both tonic and clonic phases, or either tonic or clonic phases. The time when tonic or clonic seizures was observed was determined visually by using video images captured simultaneously with iEEG signals. The transition time from tonic to clonic phases was excluded from the analyses.

### 2.5. Low- and high-frequency activity

To extract δ, θ, and HFA signals, we used bandpass filters of 2–4 Hz, 4–8 Hz, and 80–250 Hz, respectively. A bandpass device with a two-way, least-square, finite-impulse response filter (pop_eegfiltnew.m from the EEGLAB toolbox, https://sccn.ucsd.edu/eeglab/index.php) was applied to the 60-minute iEEG signals. The filter order was automatically set using the function pop_eegfiltnew.m from the EEGLAB toolbox. We calculated the amplitude of the HFA, which was an envelope of 80–250-Hz signals, in combination with the Hilbert transformation. The power of the HFA was the square of the HFA amplitude.

### 2.6. PAC analyses

The SI (Cohen, 2008) was used to measure the strength of PAC between the HFA amplitude and δ or θ phase. Hilbert transformation was performed on the bandpass-filtered signals to obtain complex-valued analytic signals [Z(t)]. The amplitude [A(t)] and phase [φ(t)] were calculated from the complex-valued signals by using Equation 1:

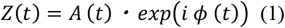

The δ and θ phases were calculated using the angle of the Hilbert transformation in the 2–4 Hz and 4–8 Hz bandpass-filtered signals, respectively. The HFA power amplitude was calculated using the squared magnitude of the envelope of the Hilbert transformation in the 80–250 Hz bandpass-filtered signal. The HFA power amplitude time series was normalized, and the phase of the amplitude was computed using the Hilbert transformation. The SI was calculated using Equation 2:

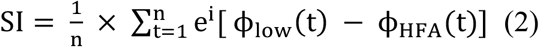

where *n* is the number of data points. We used a 1-second time window to calculate the SI in combination with the 1000 sampling rate. Therefore, in this study, n was set to 1000. The SI is a complex number; therefore, we used the magnitude of the SI, referred to as SIm. The SIm varies between 0 and 1, with 0 indicating completely desynchronized phases and 1 indicating perfectly synchronized phases.

The preferred phase of synchronization (SIp) was calculated using arctan (image [SI] / real [SI]). The SIp varies between −180° and +180°. SIm and SIp values were calculated repeatedly by shifting the 1-second time window every 33 ms, and their time-series data were obtained.

### 2.7. Bootstrapped technique and family-wise error-corrected threshold

For the statistical assessment of SIm, we shifted the phase-time series of the HFA amplitude and calculated the bootstrapped SIm (SImb) by using the δ or θ phase. This procedure was repeated 1000 times to create the distribution of SImb (Cohen, 2008), which represented the surrogate data. The maximum values of the distribution of SImb were stored at each surrogate data point, and the distribution of the maximum values was obtained. The values at 95% of the distribution of the maximum were defined as family wise error (FWE)-corrected threshold, and we applied this threshold to the observed SIm for solving multiple comparisons (Cohen, 2014). SIm values above the FWE-corrected thresholds were considered statistically significant.

### 2.8. Correlation analysis

Using all the implanted contacts in all seizures, we calculated the Spearman correlation coefficients between the normalized SIm and normalized HFA power. Values greater than +3 standard deviation (SD) or less than −3 SD were excluded as outliers. A total of 745 contacts in the tonic phase and 820 contacts in the clonic phase were used. Using the Monte Carlo method, we obtained the threshold of the correlation coefficients that achieved 80% statistical power.

### 2.9. Phase-conditioned analysis

We calculated the mean vector and performed the Rayleigh test to evaluate the nonuniformity of SIp using the CircStat toolbox (Berens, 2009). To identify the lower frequency phase to which the HFA power was coupled, we computed the average oscillation of the δ and θ bands and the average normalized HFA power. The phases of the δ and θ bands were divided into 12 intervals of 30° without overlaps: −150° ± 15°, −120° ± 15°, …, 150° ± 15°, and 180° ± 15°, resulting in 12 phase bins. The normalized HFA power was averaged for each phase bin. The values of SIp, lower frequency oscillations, and HFA power, which were obtained for significant SIm values, were used for phase-conditioned analyses.

### 2.10. Classification

We implemented a classification to distinguish between the tonic and clonic phases using δ-HFA SIm, θ-HFA SIm, and normalized HFA power. We set the varying threshold and performed binary classification. We obtained a confusion matrix, from which we calculated the sensitivity and specificity. Receiver-operating characteristic (ROC) analysis and the area under the curve (AUC) were used to compare the performance of different classifiers.

### 2.11. Statistics

For non-parametric and paired comparisons between two groups, we used the Wilcoxon signed-rank test. The AUC was also compared using the same test. The results were corrected using the Bonferroni correction for multiple comparisons.

### 2.12. Data availability

All data that were generated by or analyzed in this study are available from the corresponding authors upon reasonable request and after additional ethics approval regarding the data provision to individual institutions.

## 3. Results

### 3.1. Characteristics of investigated seizures

We enrolled six patients (five male and one female; age: 15–47 years of age, mean ± SD 24.8 ± 12.0 years) who experienced 11 focal impaired-awareness seizures (FIAS) (Table 1). All FIASs were also FBTCS, and included both tonic (10/11 seizures [91%]) and clonic (11/11 seizures, 100%) phases. The average time of the tonic phases was 22.60 ± 6.2 s and that of clonic phases was 24.82 ± 9.5 s, with no significant time-differences between tonic and clonic phases (*p* = 0.93, two-tailed Wilcoxon signed-rank test).

### 3.2. Representative tonic and clonic seizures

Results from seizure 1 (S1) in patient 1 (P1) are shown in Fig.1. S1 is also shown in the supplementary movie.

**Figure 1.**
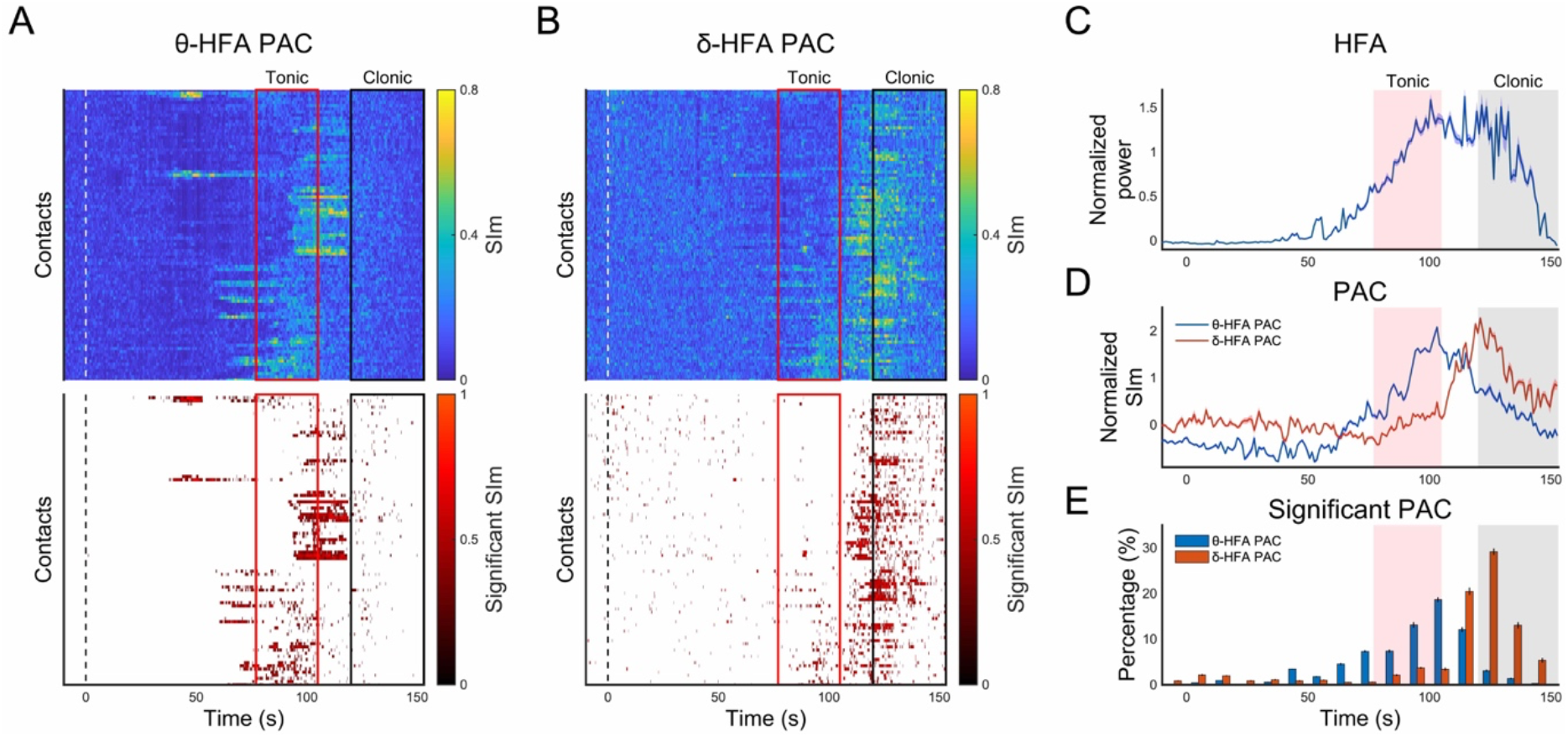
Profiles of the magnitude of the synchronization index (SIm) and high-frequency activity (HFA) related to seizure 1 (S1) in patient 1 (P1). The images were generated from a representative tonic-clonic seizure (S1-P1). Seizure onset (SO) corresponded to 0 s. The tonic and clonic phases are indicated by red square or red mesh and black square or gray mesh, respectively. Sequential SIm changes after SO are shown as topographies. Each data point of the vertical axes (A and B) corresponds to a contact. Contact names are omitted. The upper topographies indicate the raw SIm values. In the lower topographies, only significantly high SIm values over the family-wise error– corrected threshold are shown. A. The results of θ (4-8 Hz)-HFA SIm are shown. B. The results of δ (2-4 Hz)-HFA SIm are shown. C. Normalized HFA (80–250 Hz) power values were averaged across all implanted contacts. D. The normalized SIm values were averaged across all implanted contacts. E. The percentage at which significant changes occurred was calculated by dividing the number of contacts that indicated significantly high SIm values by that of all contacts.

With regard to θ-HFA PAC, more contacts showed significantly higher θ-HFA SIm values in the tonic than in the clonic phase (Fig. 1A). In contrast, δ-HFA PAC showed more contacts, indicating significantly higher δ-HFA SIm values, in the clonic than in the tonic phase (Fig. 1B). The average values of the normalized HFA power and normalized θ-HFA SIm across all contacts peaked in the late tonic phase (Fig. 1C, 1D); however, the average values of normalized δ-HFA SIm across all contacts reached a maximum in the early clonic phase (Fig. 1D). Similarly, the percentage of contacts indicating significantly high SIm values showed that θ-HFA PAC peaked in the late tonic phase and that δ-HFA PAC peaked in the early clonic phase (Fig. 1E).

Figure 2 shows the time series of raw iEEG signals, HFA power, and lower frequency oscillations obtained from one contact, which indicated significantly high SIm values. These time series were observed upon the appearance of significantly high θ-HFA SIm values in the tonic phase and significantly high δ-HFA SIm values in the clonic phase. In both the tonic and clonic phases, the HFA power burst occurred at the peak of the iEEG spike (red mesh areas in Fig. 2). In the tonic phase, it seemed that the HFA power burst occurred at the baseline (0 μV) between the peak and trough of the θ oscillations (Fig. 2A). In contrast, the HFA power burst in the clonic phase tended to occur at the trough of δ oscillations (Fig. 2B). We inferred that during coupling occurrence, the HFA power would couple with certain phases of the lower frequency band, and the main phase coupled with the HFA power would be different between θ-HFA PAC in the tonic phase and δ-HFA PAC in the clonic phase. Therefore, we performed phase-base analyses (see Section 3.5.)

**Figure 2.**
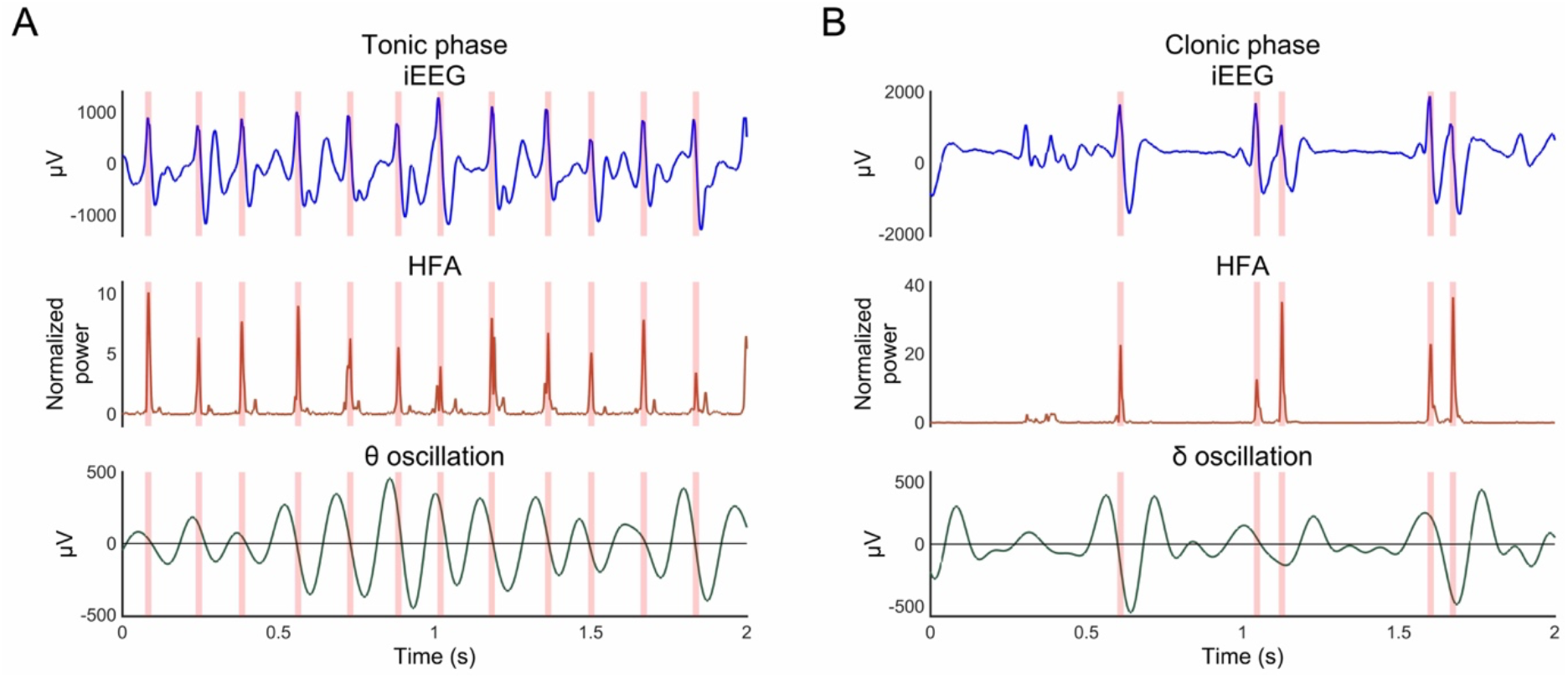
Intracranial electroencephalogram (iEEG) signals, high-frequency activity (HFA) power, and lower-frequency oscillations. Upper column: iEEG signals. Middle column: normalized HFA (80–250 Hz) power time series. Lower column: θ (4-8 Hz) oscillations (A) or δ (2-4 Hz) oscillations (B). These time series were calculated from one contact, which indicated significantly high θ-HFA magnitude of synchronization index (SIm) values in the tonic phase (A) or significantly high δ-HFA SIm values in the clonic phase (B). The time points of the iEEG spike were indicated by red mesh areas.

### 3.3. Profiles of θ-HFA PAC and δ-HFA PAC

We calculated the normalized SIm and normalized HFA power across all contacts, indicating significantly high SIm values over the FWE-corrected threshold within all seizures, and compared them between the tonic and clonic phases (Fig. 3). In θ-HFA PAC, the normalized θ-HFA SIm values were significantly higher in the tonic than in the clonic phase (*p* = 0.0039, two-tailed Wilcoxon signed-rank test); however, significantly higher normalized δ-HFA SIm values were observed in the clonic than in the tonic phase (*p* = 0.0020, two-tailed Wilcoxon signed-rank test) (Fig. 3A). The normalized HFA power during the occurrence of significant SIm was significantly higher in the tonic than in the clonic phase related to both θ-HFA PAC (*p* = 0.014, two-tailed Wilcoxon signed-rank test) and δ-HFA PAC (*p* = 0.049, two-tailed Wilcoxon signed-rank test) (Fig. 3B).

**Figure 3.**
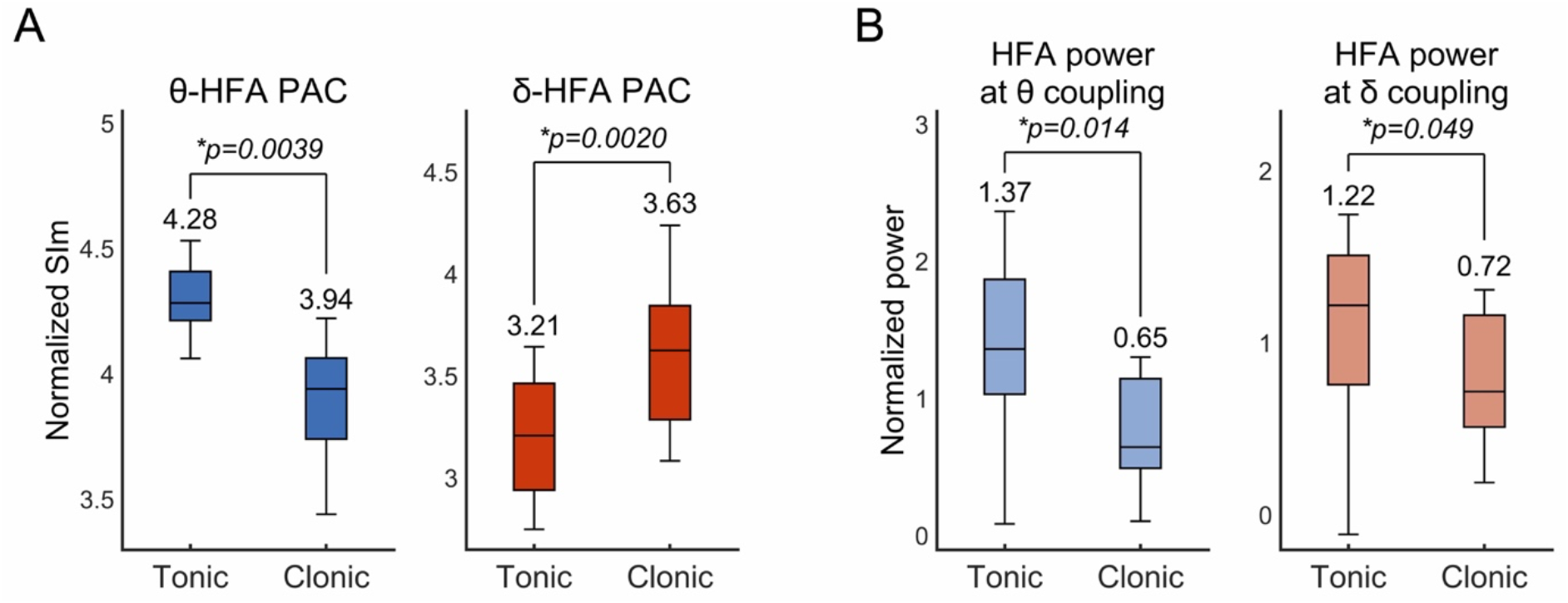
The profiles of high-frequency activity (HFA) and phase-amplitude coupling (PAC) in tonic and clonic phases. The results calculated from contacts indicating a significantly high magnitude of synchronization index (SIm) are shown as box-and-whisker plots, in which the median values for each group are displayed. A. Normalized SIm of θ-HFA PAC was significantly higher in the tonic than in the clonic phase. However, this was reversed in δ-HFA PAC. B. The normalized HFA power was calculated using contacts indicating a significantly high SIm of θ-HFA PAC or δ-HFA PAC. The normalized HFA power in the tonic phase was significantly higher than that in the clonic phase for both θ-HFA PAC and δ-HFA PAC. We used the Wilcoxon signed-rank test for the statistical evaluation.

We evaluated the percentage of significant SIm values across all contacts within all seizures (Fig. 4). In the late tonic phase, we observed the maximum percentage related to θ-HFA PAC (median, 17.68%), which was significantly higher than that in the other three stages including the early tonic, early clonic, and late clonic (corrected *p* = 0.041 between the early tonic and late tonic, 0.041 between the late tonic and early clonic, 0.0059 between the late tonic and late clonic; two-tailed Wilcoxon signed-rank test with Bonferroni correction) (Fig. 4A). In the early clonic phase, we observed the maximum percentage related to δ-HFA PAC (median, 18.06%), which was significantly higher than that in tonic phases (corrected *p* = 0.0059 between the early tonic and early clonic, 0.029 between the late tonic and early clonic; twotailed Wilcoxon signed-rank test with Bonferroni correction) (Fig. 4B). We concluded that the HFA power and θ-HFA PAC occur more obviously in the tonic than in the clonic phase and that δ-HFA PAC occurs more obviously in the clonic than in the tonic phase.

**Figure 4.**
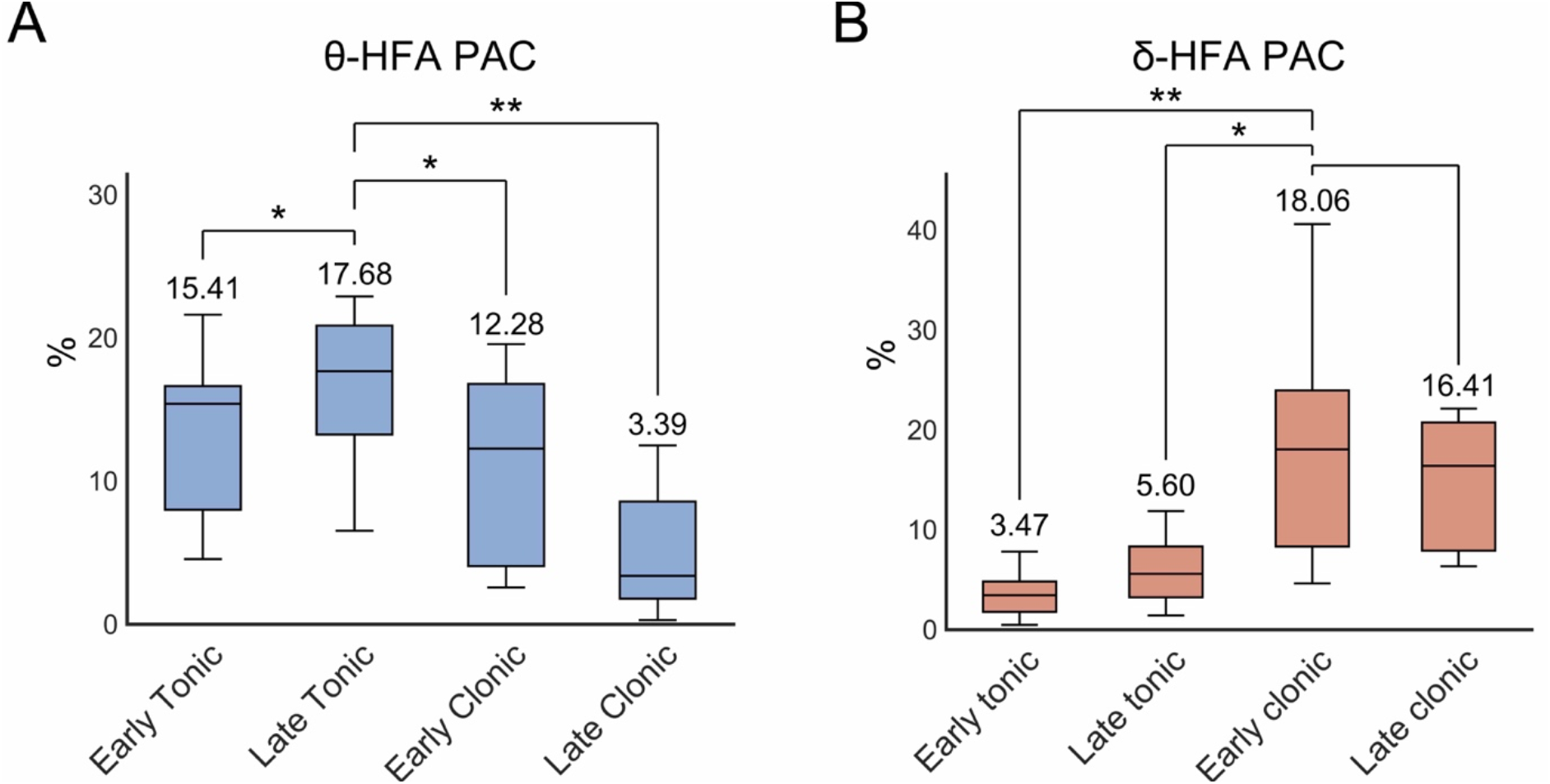
Percentages at which significant magnitude of synchronization index (SIm) values occur. The percentages at which significantly high SIm occur were evaluated at four stages: early or late and tonic or clonic phases. The results are shown as box-and-whisker plots, in which the median values at each stage are displayed. A. The percentage of significantly high θ-high-frequency activity (HFA) phase-amplitude coupling (PAC) peaked at the late tonic phase. B. The percentage of significantly high δ-HFA PAC peaked in the early clonic phase. *corrected p < 0.05, ** corrected p < 0.01, Wilcoxon signed-rank test, three multiple comparisons corrected by Bonferroni method.

### 3.4. Correlation between normalized HFA power and normalized SIm

By using all implanted contacts, we calculated the correlation coefficients (*r*) in combination with the normalized HFA power and θ-HFA or δ-HFA normalized SIm (Fig. 5). We observed a stronger positive correlation related to θ-HFA PAC in the tonic than in the clonic phase. The *r* values related to θ-HFA PAC decreased from the tonic phase (*r* = 0.39) to the clonic phase (*r* = 0.19) (Fig. 5A). In contrast, the *r* values related to δ-HFA PAC increased from the tonic (*r* = 0.11) to the clonic phase (*r* = 0.29) (Fig. 5B). We set the threshold of the correlation coefficients that achieved 80% statistical power, which was ± 0.1. Therefore, our findings demonstrate a significant positive correlation.

**Figure 5.**
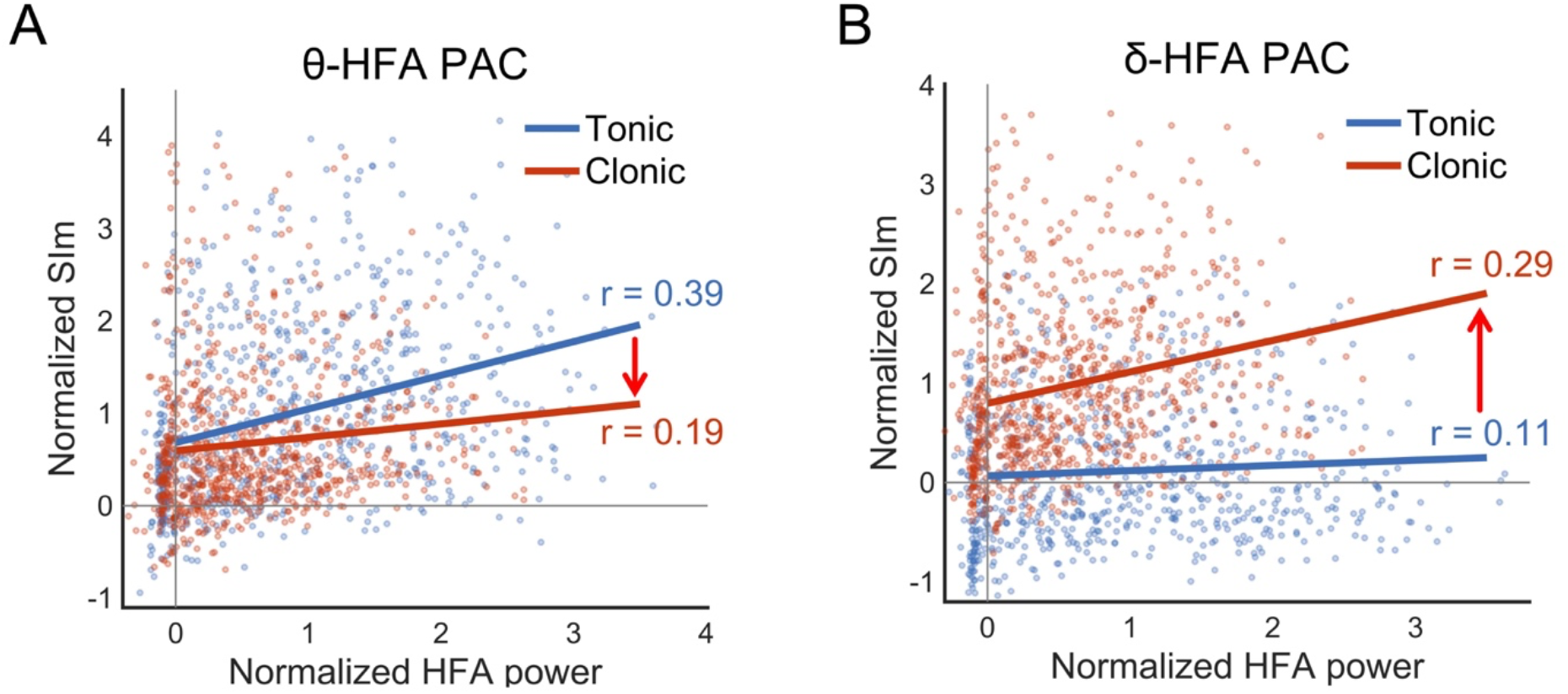
Correlation between normalized high-frequency activity (HFA) and normalized magnitude of synchronization index (SIm). Using all implanted contacts, correlation coefficients (*r*) were calculated between the normalized HFA and normalized SIm. A. In θ-HFA phase-amplitude coupling (PAC), a stronger positive correlation was observed in the tonic than in the clonic phase. B. In δ-HFA PAC, a stronger positive correlation was observed in the clonic than in the tonic phase.

### 3.5. Phase-based analyses

Phase-based analyses were used to investigate the differences between θ-HFA PAC in the tonic phase and δ-HFA PAC in the clonic phase. We calculated the mean vectors of SIp that were obtained when SIm was significantly higher than the FWE-corrected threshold (Fig. 6A). When θ-HFA SIm was significantly high in the tonic phase, the SIp values were unevenly distributed at approximate 75°–90°. The angle of the mean vector was 73.09°, and significant non-uniformity was observed (*p* = 0, Rayleigh test; left panel in Fig. 6A). When δ-HFA SIm was significantly high in the clonic phase, the angle of the mean vector was 33.90°, and significant non-uniformity was observed (*p* = 0, Rayleigh test). However, there were two large groups of δ-HFA SIp values at approximately 120° and 330° (right panel in Fig. 6A).

**Figure 6.**
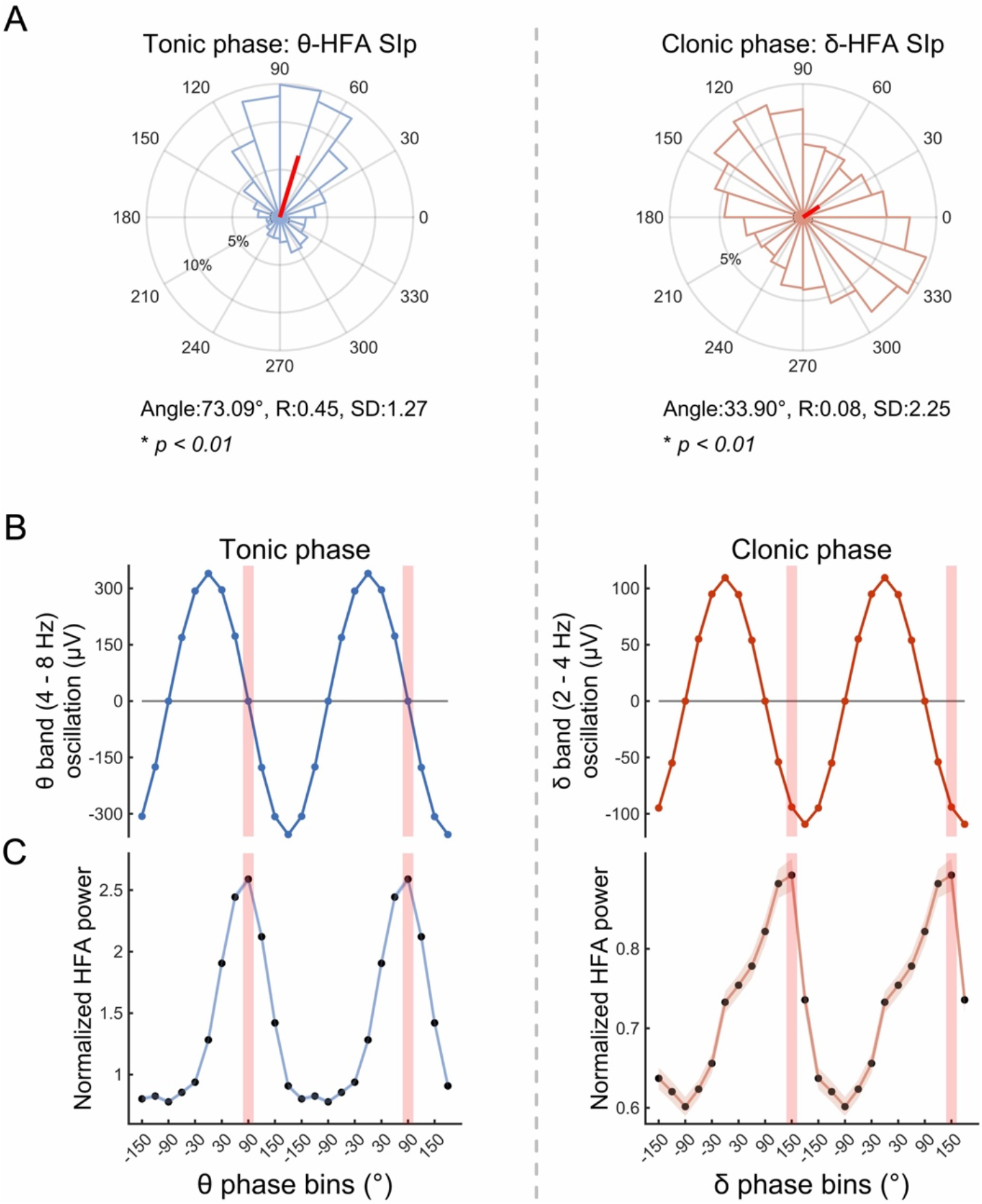
Phase-based analyses of phase-amplitude coupling (PAC). Phase-based analyses were performed using values obtained when statistically high magnitude of synchronization index (SIm) values were observed. Left panels indicate the results related to θ-HFA PAC in the tonic phase, and right panels indicate the results related to δ-HFA PAC in the clonic phase. A. We calculated the average values of the preferred phase of synchronization (SIp). The angle, length (R), and standard deviation (SD) of the mean vector (red bold lines) are indicated. The p-values were calculated using the Rayleigh test. The θ or δ oscillations of the θ or δ phase (B) and normalized high-frequency activity (HFA) power tuned by the θ or δ phase (C) are displayed. The phases at which the normalized HFA power peaked were indicated by red mesh areas. The error bars in (B) and (C) indicate the 95% confidence intervals. In (B), the error bars are too narrow, so they are not visible.

Next, phase-tuned oscillations and the normalized HFA power were calculated when the SIm became statistically high. Figure 6B depicts θ oscillations tuned by the phases of the θ band in the tonic phase (left panel) and δ oscillations tuned by the phases of the δ band in the clonic phase (right panel). Figure 6C depicts the normalized HFA power tuned by the phases of the θ band in the tonic phase (left panel) and that tuned by the phases of the δ band in the clonic phase (right panel). When θ-HFA SIm was significantly high in the tonic phase, the normalized HFA power peaked at the baseline (0 μV/90°) between the peak and trough of the θ oscillations (left panels in Fig. 6B, C, red mesh areas). In contrast, when δ-HFA SIm values were statistically high in the clonic phase, the normalized HFA power peaked at around the trough (150°) of the δ oscillation (right panels in Fig. 6B, C, red mesh areas). These results were concordant with the individual results shown in Figure 2, indicating that the main phase of the δ band in the clonic phase is approximately 120°–150° and not 330°.

### 3.6 Classification

We evaluated whether the HFA power, θ-HFA SIm, and δ-HFA SIm accurately discriminated between the tonic and clonic phases (Fig. 7). We found that the AUC of δ-HFA SIm was at its maximum and significantly higher than that of the HFA power and that of θ-HFA SIm (corrected *p* = 6.31 × 10^-164^ for HFA power vs δ-HFA SIm, corrected *p* = 9.67 × 10^-165^ in θ-HFA SIm vs δ-HFA SIm, two-tailed Wilcoxon signed-rank test, two multiple comparisons corrected by Bonferroni method).

**Figure 7.**
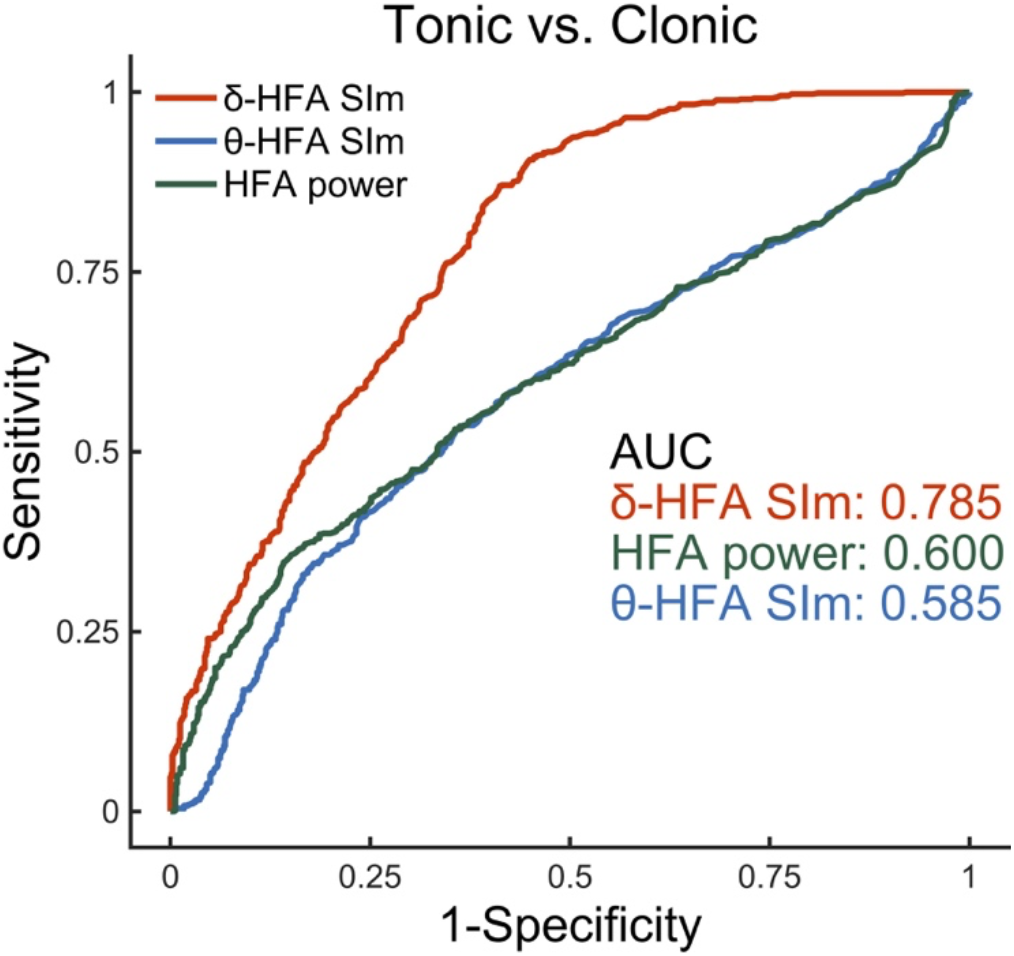
Receiver operating characteristic curves The tonic and clonic phases were classified using normalized high-frequency activity (HFA) power, θ-HFA magnitude of the synchronization index (SIm), and δ-HFA SIm. The values of area under the curve (AUC) were indicated.

## 4. Discussion

In this study, we hypothesized that θ-HFA coupling occurs in the tonic phase of FBTCS and that δ-HFA coupling occurs in the clonic phase of FBTCS. By extracting HFA using a 80–250-Hz bandpass filter, δ band by a 2–4 Hz bandpass filter, and θ band by a 4–8 Hz bandpass filter, we demonstrated that the phase of the θ band is coupled with the HFA power in the tonic phase, whereas that of the δ band is coupled with HFA power in the clonic phase. In the tonic phase, the HFA power peaked at the baseline of θ oscillations, whereas, in the clonic phase, it peaked at the trough of δ oscillations. The PAC between the δ phase and HFA power amplitude discriminates between the clonic and tonic phases more significantly than does the HFA power only and θ-HFA PAC. Therefore, we inferred that δ-HFA coupling might reflect neurophysiological features that are specific to tonic seizures.

Local ictal discharges propagate bilaterally through the corpus callosum, which induces tonic-clonic phases (Brodovskaya and Kapur, 2019, Wieshmann et al., 2015). Abnormally increased activity in subcortical structures such as the cerebellum, basal ganglia, brainstem, and thalamus, along with decreased activity in the association cortex, may play a crucial role in FBTCS (Blumenfeld et al., 2009). FBTCS was previously suspected to reflect a broken balance between basal ganglia inhibition and thalamic synchronization (He et al., 2020). It is understood by the propagation model and its relevance to the thalamus and basal ganglia, and some studies have shown disrupted whole-brain network interactions between different brain areas in the pathophysiology of FBTCS (Sinha et al., 2021). Although our results regarding θ-HFA coupling in the tonic phase and δ-HFA coupling in the clonic phase were obtained from measuring cortical activities, we inferred that these θ and δ activities might reflect the neurophysiological features of subcortical areas.

Previous studies have shown that ictal-HFA at the SOZ is coupled with the θ phase (Hashimoto et al., 2021b, Ibrahim et al., 2014), and that θ-HFA coupling discriminates normal brain regions from the SOZ (Amiri et al., 2019). Our results additionally showed that propagated HFA changes in the tonic phase, which were obtained not only from the SOZ but also from the seizure propagation zone, were also coupled with the θ phase. Animal experiments have shown that, in the epileptic brain, the hippocampal θ rhythm is increased (Kitchigina and Butuzova, 2009). Thus, the θ band may be the key frequency band representing the neurophysiological processing involved in both seizure generation and propagation.

Coupling between HFA and δ bands has been investigated in previous studies associated with epileptic spasm (Nariai et al., 2011) or an interictal state (Amiri et al., 2016). In our previous study, we showed the time lag by which θ-HFA PAC precedes δ-HFA PAC in ictal states (Hashimoto et al., 2021b). Another study also showed a shift from to 4–5 Hz (θ band) to 1–2 Hz (δ band), which was coupled with HFA throughout seizures (Grigorovsky et al., 2020). In this study, we confirmed that δ-HFA PAC occurred mainly in the clonic phase of FBTCS. Previous studies have demonstrated that in post-ictal states, increased δ power is observed more in FBTCS (Yang et al., 2012), and that δ-γ (30-50 Hz) PAC could be a biomarker of post-ictal generalized EEG suppression. Although previous studies have implied that δ band activities are mainly involved in post-ictal states, using ROC analyses, we additionally showed that δ-HFA PAC may be a potential biomarker for discriminating clonic from tonic phases in FBTCS.

The amygdala is involved in epileptogenesis, and the duration of burst discharges sets the frequency of network oscillation, which is in the δ band in both normal and seizure conditions (Chou et al., 2020). Moreover, the frequency of epileptic EEG activities has been shown to be involved in the thalamocortical network (Dichter, 1997). Therefore, we inferred that the δ band is the key frequency band in the clonic phases, which is the late stage of FBTCS to post-ictal states, and that the amygdala and thalamus are involved in the generation of the δ rhythm.

PAC methods have been used to investigate the relationship between high- and low-frequency activities (Cohen, 2008). PAC has been observed in both physiological and pathological neural processing. The former includes hand movement (Yanagisawa et al., 2012), swallowing (Hashimoto et al., 2021c) and somatosensory processing (Lakatos et al., 2008), and the latter includes epileptic seizures (Hashimoto et al., 2020b, 2021b, Iimura et al., 2018), and Parkinson’s disease (de Hemptinne et al., 2015). It was reported that ictal HFA amplitudes are coupled with δ (Iimura et al., 2018, Nariai et al., 2011), θ (Hashimoto et al., 2021b, Ibrahim et al., 2014), and α (Ibrahim et al., 2014) phases, while β-HFA coupling has been reported as a useful marker for seizure detection (Edakawa et al., 2016). Moreover, coupling θ waves with HFOs has been reported to effectively discriminate normal brain regions from the SOZ (Amiri et al., 2019). PAC is an effective methodology for seizure analyses. Using this method enabled us to demonstrate that the main low-frequency band coupled with HFA amplitude varies between the tonic and clonic phases.

The HFA power is strongly correlated with the neural firing rate (Ray et al., 2008), and, in animal studies, neural spiking was found to be locked to the trough of α oscillations (Haegens et al., 2011). In a previous study of physiological PAC, high gamma amplitude during high PAC values peaked around the trough of α oscillations (Hashimoto et al., 2021c, Yanagisawa et al., 2012). Moreover, in a study of pathological PAC related to epilepsy, the HFA amplitude during high PAC was time-locked to the trough of lower-frequency oscillations (Ibrahim et al., 2014). During the clonic phase, the HFA amplitude was tuned to the trough of δ oscillations, which is concordant with the findings of previous reports. In our previous study, the HFA amplitude at the SOZ was tuned at the trough of the θ band (Hashimoto et al., 2021b). In this study, the HFA amplitude in the tonic phase was tuned at the baseline of the θ band. Since apart from the SOZ we also focused on the seizure propagation zone, this difference in phase, which tuned the HFA amplitude, might reflect the difference between the SOZ and seizure propagation zone.

HFAs have been observed during seizures (Akiyama et al., 2011, Ochi et al., 2007), and HFOs, a subgroup of HFAs (Ayoubian et al., 2013), have been suggested as useful biomarkers for the detection of the SOZ (Wu et al., 2014). HFOs are reported to be superimposed on spikes (Wang et al., 2013), in agreement with our finding of a superimposed HFA power on a spike (Figure 2). Previous studies have shown that prior to the onset of bilateral tonic-clonic movements, the ripple (80-250 Hz) density in the SOZ is higher in FBTCS than in focal seizures (Schönberger et al., 2019). We additionally demonstrated that, in FBTCS, the HFA power is significantly higher in the tonic than in the clonic phase. Although the HFA power decreased in the clonic phase, as compared with its value in the tonic phase, δ-HFA PAC and correlation coefficients between SIm and HFA power increased. Therefore, we inferred that the neurophysiological characteristic band specific to the clonic phase might be the δ band.

This study has some limitations. First, the sample size was small. Therefore, we evaluated all seizures without adjusting for the number of seizures in each patient. Second, all seizures were recorded after extensively reducing antiepileptic drugs; thus, they might not represent the patients’ habitual seizures. However, we consider that our analysis is independent of this issue because reducing the medication did not affect the morphology of discharges at onset or the duration of the contralateral spread (So and Gotman, 1990). Finally, because this study included only patients with focal epilepsy, our findings may not be generalizable to patients with generalized epilepsy.

## 5. Conclusions

By analyzing FBTCS using PAC, we revealed that the θ band is the main low-frequency band modulating the HFA power during the tonic phase, while the δ band modulates the HFA power during the clonic phase. Moreover, ROC curve analysis indicated that δ-HFA PAC discriminated well between the tonic and clonic phases. We conclude that the θ and δ oscillations represent the characteristic neurophysiological processes involved in the tonic and clonic phases, respectively.

## Funding

This study was supported by a grant from Grants-in-Aid for Early-Career Scientists (KAKENHI; grant nos. JP18K18366 (Hiroaki Hashimoto), JP21K16629 (Hiroaki Hashimoto)), which is funded by the Japan Society for the Promotion of Science (JSPS; Tokyo, Japan).

## Declaration of interest

None.

## Abbreviations

AUC: area under the curve
EEG: electroencephalogram
FBTCS: focal to bilateral tonic-clonic seizures
FIAS: focal impaired awareness seizure
FWE: family-wise error
HFA: high-frequency activity
HFO: high-frequency oscillation
iEEG: intracranial electroencephalogram
MRI: magnetic resonance imaging
PAC: phase-amplitude coupling
ROC: receiver-operating characteristic
SD: standard deviation
SI: synchronization index
SIm: magnitude of SI
SImb: bootstrapped SIm
SIp: preferred phase of synchronization
SOZ: seizure onset zone
3D: three-dimensional

## Author Contributions

H.H. conceived the study, collected the data, created the MATLAB program, analyzed the data, created all figures and the movie, and was primarily responsible for writing the manuscript. H.M.K., N.T., S.O., M.H., and H.K. performed the epileptic surgery. All authors clinically cared for and evaluated the patients. M.H. and H.K. supervised the study. All authors have reviewed the manuscript.

## Movie

Multimodal data obtained from patient 1-seizure 1 are shown; normalized high-frequency activity (HFA) (80–250 Hz) power distribution map (A), θ (4-8 Hz)-HFA phase-amplitude coupling (PAC) (magnitude of synchronization index: SIm) distribution map (B), δ (2-4 Hz)-HFA PAC (SIm) distribution map (C), and intracranial electroencephalogram (iEEG) signals (D). The movie contains the results of the tonic and clonic phases. Circles in A, B, and C correspond to implanted intracranial electrode contacts.

In the HFA power distribution map (A), the red circles were scaled linearly with the normalized HFA power changes. Fluctuations in red circles were observed at a faster rhythm in the tonic than in the clonic phase.

We display only statistically significant SIm values to which the family-wise error– corrected threshold was applied. Significantly high SIm values were scaled from black to red. In the tonic phase, we observed more contacts with significantly high θ-HFA SIm values rather than significantly high δ-HFA SIm values. In the clonic phase, more contacts showed significant SIm values in δ-HFA PAC than in θ-HFA PAC.

Only some representative contacts that indicated typical iEEG changes are shown on the vertical axes in D.

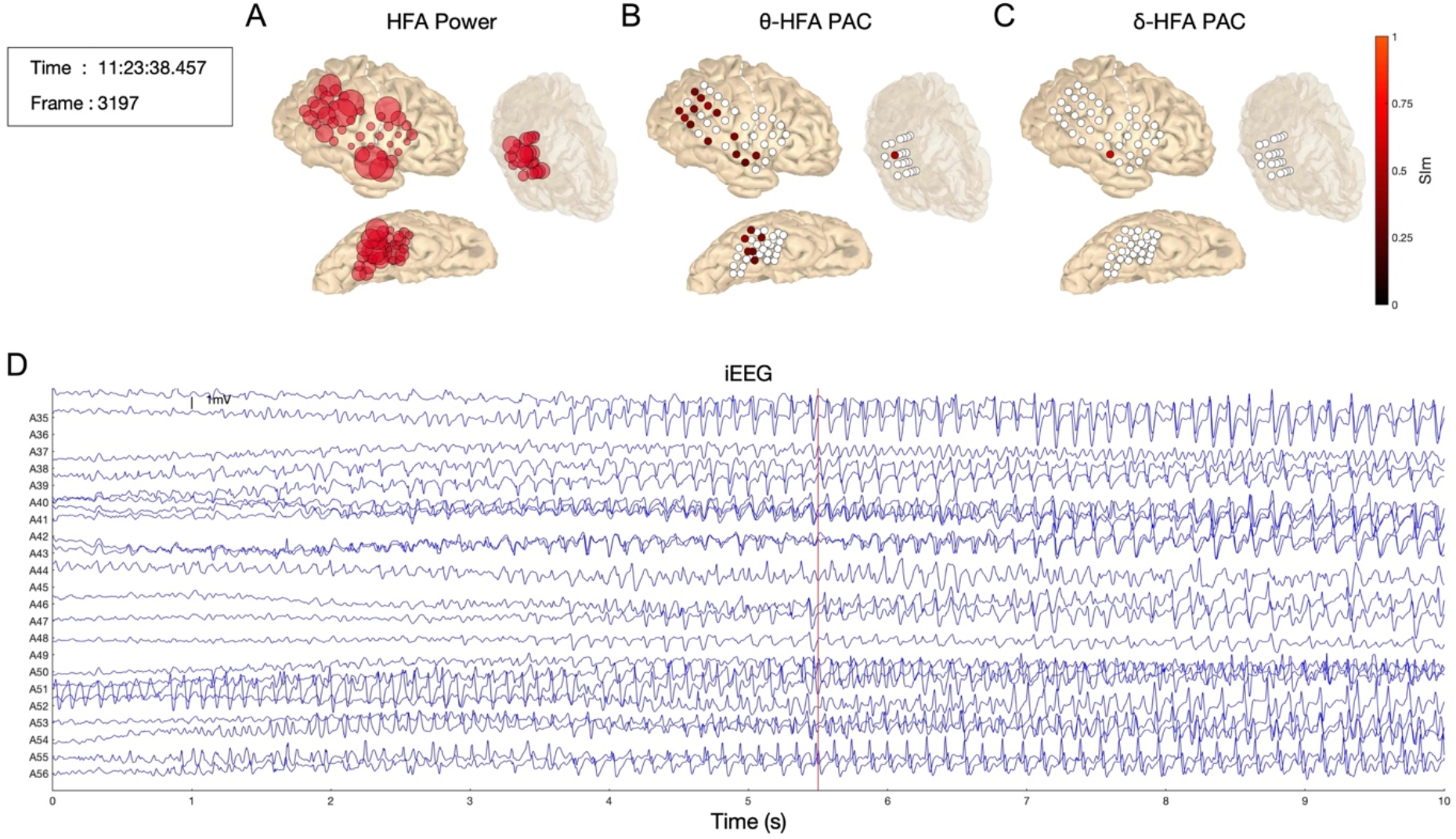

## Notes

### Competing Interest Statement

The authors have declared no competing interest.

